# Apoptosis recognition receptors regulate skin tissue repair in mice

**DOI:** 10.1101/2023.01.17.523241

**Authors:** Olivia Justynski, Kate Bridges, Will Krause, Maria Fernanda Forni, Quan Phan, Teresa Sandoval-Schaefer, Ryan Driskell, Kathryn Miller-Jensen, Valerie Horsley

## Abstract

Apoptosis and clearance of apoptotic cells via efferocytosis are evolutionarily conserved processes that drive tissue repair. However, the mechanisms by which recognition and clearance of apoptotic cells regulate repair are not fully understood. Here, we use single-cell RNA sequencing to provide a map of the cellular dynamics during early inflammation in mouse skin wounds. We find that apoptotic pathways and efferocytosis receptors are elevated in fibroblasts and immune cells, including resident Lyve1^+^ macrophages, during inflammation. Interestingly, human diabetic foot wounds upregulate mRNAs for apoptotic genes and display increased and altered efferocytosis signaling via the receptor Axl. During early inflammation in mouse wounds, we detect upregulation of Axl in dendritic cells and fibroblasts via TLR3-independent mechanisms. Inhibition studies *in vivo* in mice reveal that Axl signaling is required for wound repair but is dispensable for efferocytosis. By contrast, inhibition of another efferocytosis receptor, Timd4, in mouse wounds decreases efferocytosis and abrogates wound repair. These data highlight the distinct mechanisms by which apoptotic cell detection coordinates tissue repair and provides potential therapeutic targets for chronic wounds in diabetic patients.

## Introduction

Proper tissue function and homeostasis require efficient and effective repair of injury. Repair of mammalian tissues requires highly dynamic changes in cellular heterogeneity and communication to properly heal tissue, usually resulting in a scar rather than true tissue regeneration. Cell death is a common event during tissue injury, and several studies from hydra to mice have shown the importance of apoptosis in the initiation of inflammation to drive reparative processes (Greenhalgh 1998). Proper initiation and subsequent resolution of inflammation is essential for tissue repair and progression to the proliferation stage of healing, when fibroblasts, blood vessels, and other tissue specific cells proliferate and migrate, forming new tissue to repair the wound. While several signaling factors have been shown to induce apoptosis in wounds (Guerin et al. 2021), the mechanisms by which apoptotic cells are recognized and regulate tissue repair are not well understood.

The skin is an excellent model to define the mechanisms by which apoptotic cells regulate tissue repair. After injury, mammalian skin undergoes stages of repair including inflammation, which removes debris and pathogens. As inflammation regresses, the proliferative phase leads to the coordination of epidermal keratinocytes, fibroblasts, endothelial, and immune cells to reseal the epidermal barrier and generate a reparative scar (Eming et al. 2014). Apoptosis occurs after skin injury and phagocytosis of apoptotic cells – or efferocytosis – by macrophages reduces inflammatory signaling and repair in several tissues (Bosurgi et al 2017a, Peiseler & Kubes 2019). Yet, it is unclear how apoptosis controls skin wound healing.

Apoptotic cell death is characterized by cytomorphological alterations, DNA fragmentation, activation of caspases and other regulators, and finally membrane alterations including outer membrane exposure of phosphatidylserine (PtdSer), which allows the recognition of apoptotic cells by cellular receptors on phagocytes (Elmore 2007). The most well-studied receptors that allow phagocytes to bind and phagocytose apoptotic cells include the TAM (Tyro3, Axl, and Mertk) tyrosine kinases and the TIM (T cell immunoglobulin and mucin domain) family of receptors (Lemke 2019). While TIM receptors can directly bind PtdSer, TAM receptors require their ligands growth-arrest-specific 6 (Gas6) and protein S (Pros1) to bind to PtdSer (Lew et al. 2014, Elliott et al. 2017).

To understand the cellular and molecular mechanisms by which apoptosis regulates skin wound healing, we performed single-cell RNA sequencing (scRNA-seq) on cells from murine wound beds 24 and 48 hours after injury. We found that transcriptional alterations in apoptotic pathways occur in this interval in fibroblasts, monocytes/macrophages, neutrophils, and dendritic cells. In addition, inhibition of two efferocytosis receptors, Axl and Timd4, abrogates proper wound repair. The results provide an atlas of cellular dynamics during the early stages of wound healing and reveal the essential role of the recognition and clearance of apoptotic cells in driving tissue repair after injury.

## Results

### Dramatic transcriptional heterogeneity during early skin inflammation after injury

To assess cellular and molecular heterogeneity of the wound bed during the inflammatory phase, we performed scRNA-seq on cells isolated from 4 mm full-thickness biopsy punches on mouse back skin at 24 and 48 hours (h) after injury (Shook et al. 2016). To ensure that we also captured the immediately adjacent tissue as well as cells that may have migrated into the wound site, we used a 6mm biopsy punch to collect the tissue before isolating cells using enzymatic digestion (**Fig. 1A**). By training a neural network to identify cell types based on expression of established marker genes (as in Wasko et al. 2022) (**Fig. S1A-B**), we classified 4 major cell types in the scRNA-seq data, including monocytes/macrophages (Mono/MO), neutrophils (Neut), dendritic cells (DC), and fibroblasts (FB) at both time points (**Figs. 1B-E**).

**Figure 1:**
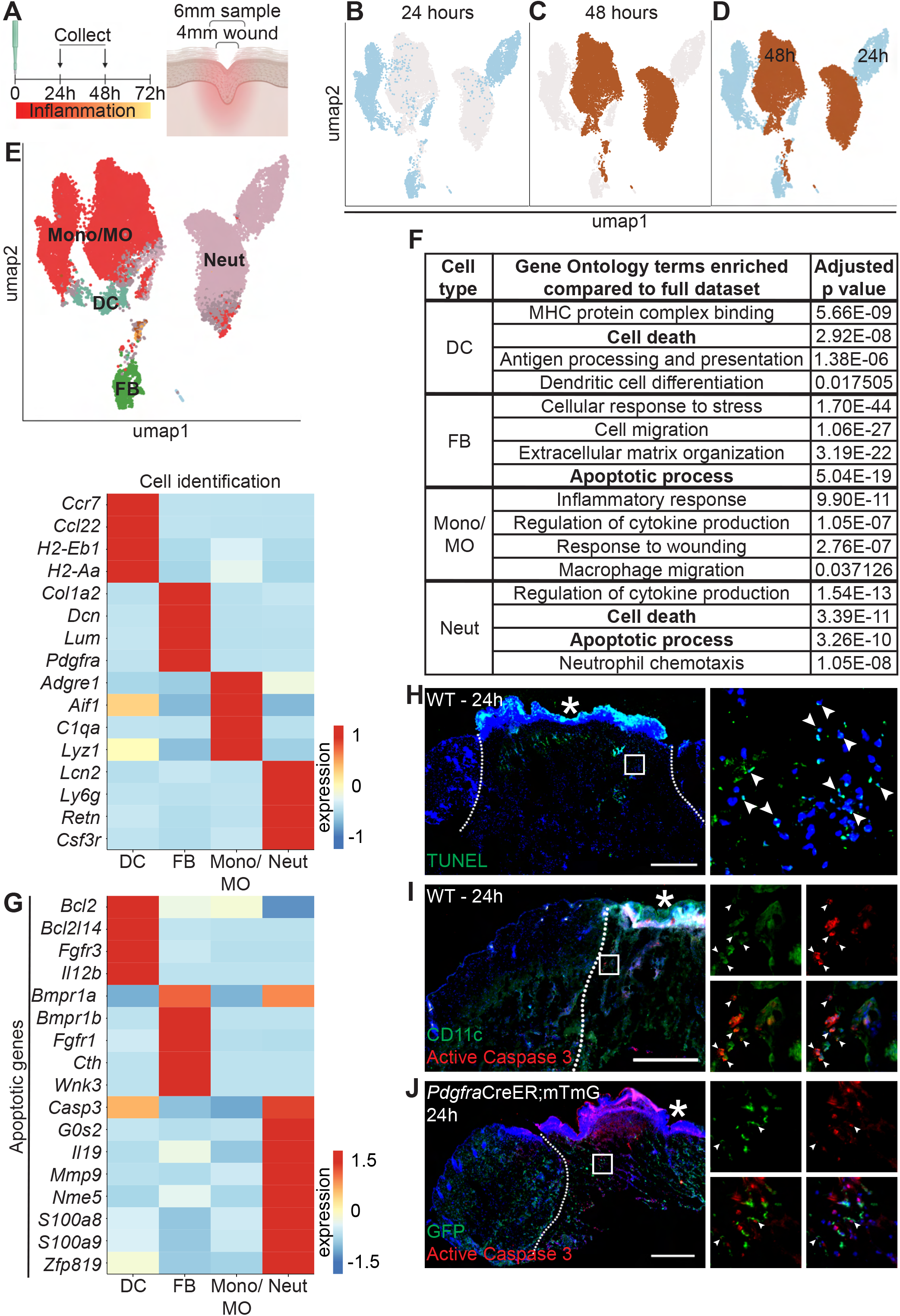
Dynamic transcriptional heterogeneity and apoptosis are observed in murine wound beds 24h and 48h after injury. A. Schematic of experimental design B. UMAP plot of scRNA-seq data for cells from 24h wound beds in murine back skin. C. UMAP plot of scRNA-seq data for cells from 48h wound beds in murine back skin. D. UMAP plot of scRNA-seq data for cells from both 24h and 48h wound beds in murine back skin annotated by timepoint. E. **Top:** UMAP plot of scRNA-seq data for cells from 24h and 48h wound beds annotated by cell identity. **Bottom:** Heatmap of differentially expressed marker genes in 24h and 48h wound beds. F. Gene ontology terms enriched in each cell type compared to the full dataset. G. Heatmap of differentially expressed apoptosis-related genes from 24h and 48h wound beds. H. Immunostaining for TUNEL (green) in wound bed 24h after injury. Arrows indicate TUNEL^+^ cells. I. Immunostaining for CD11c (green) and Active caspase 3 (red) in wound bed 24h after injury. Arrows indicate double-positive CD11c^+^ Active caspase 3^+^ cells. J. Immunostaining for GFP (green) and Active caspase 3 (red) in PdgfraCreERmTmG wound bed 24h after injury. Arrows indicate double-positive GFP^+^ Active caspase 3^+^ cells. * indicates scab. Scale bars = 500µm.

Surprisingly, the 24h and 48h samples clustered separately with minimal overlap, suggesting that dramatic changes occurred in the first two days of inflammation (**Figs. 1B-D**). The absolute number of macrophages and neutrophils increased >2 fold over this interval (**Fib. S1C**). While approximately 70% of each of the three immune cell populations was collected at 48h, the majority of fibroblasts were found in the 24h population (**Fig. S1D**). We also found that genes upregulated by each cell type were markedly different between timepoints (**Fig. S1E**), indicating major changes to expression patterns in the same cell type over time.

To explore the changes in the major cell types in early wound beds (fibroblasts, neutrophils, DCs, and monocytes/macrophages), we analyzed the genes that were significantly upregulated in each group relative to the mRNAs expressed by the full dataset to determine gene ontology terms that were enriched for each cell type (**Fig. 1F**). Monocytes/macrophages, neutrophils, and DCs upregulated mRNAs involved in their specific function in inflammation for cytokine production, chemotaxis, and antigen presentation, respectively. Similarly, fibroblasts uniquely upregulated genes involved in extracellular matrix (ECM) organization. Interestingly, DCs, fibroblasts, and neutrophils upregulated genes involved in apoptosis and cell death (**Figs. 1F and 1G**). Neutrophils upregulated the largest number of mRNAs for apoptosis including *Casp3*, which is phosphorylated to activate apoptosis, *S100a8* and *S100a9*, which induce apoptosis of several cell types, and *G0s2*, which binds Bcl2 to promote apoptosis (Yui et al. 2003, Welch et al. 2009). Fibroblasts and DCs also upregulated apoptotic genes ranging from receptors (*Fgfr1* and *Bmpr1a*), cytokines (*Il12b*), to mediators of apoptotic pathways (*Bcl2l14* and *Cth*).

Since apoptosis also involves post-transcriptional activation of several proteins, we sought to confirm *in vivo* that apoptosis occurred during early skin wound inflammation. Sections of mouse wounds at 24h (**Fig. 1H**) and 48h (**Fig. S1F**) were processed with TUNEL assay to detect double-stranded DNA breaks characteristic of apoptosis. Staining with antibodies against phosphorylated-caspase 3, its active form, confirmed the presence of CD11c^+^ dendritic cells (**Fig. 1I**) and Pdgfra^+^ fibroblasts (**Fig. 1J**) that were apoptotic in 24h wound beds. However, a robust population of apoptotic cells was not observed. We confirmed this result via flow cytometry, using the cell viability stain Propidium iodide (PI) and Annexin V (AV) staining. Annexin V binds to exposed PtdSer, thus marking apoptotic cells. We found that live cells made up the majority of single cells in non-wounded (NW) skin as well as 24h and 48h wound beds, and that no increase of apoptotic cells was observed in wound bed samples (**Figs. S1G**). These data suggest that during inflammation, multiple cell types upregulate cell death pathways and efferocytosis of apoptotic cells is a rapid and efficient process.

### Apoptosis recognition receptors, ligands, and downstream factors are expressed in the wound bed

Given the upregulation of apoptotic genes and dramatic changes in phagocytic neutrophils and macrophages we observed during early inflammation, we inspected expression levels of mRNAs for efferocytosis receptors, their ligands, and downstream factors in scRNA-seq data from 24h and 48h wounds (**Fig. 2A**). Interestingly, fibroblasts upregulated several genes encoding efferocytosis receptors (*Tyro3, Axl*, and *Itgav*), as well as genes encoding for ligands *Gas6, Pros1, C3, C4b*, and *Mfge8*. DCs, macrophages, and neutrophils also upregulated several receptors that mediate detection of apoptotic cells. Macrophages were enriched for the downstream activators *Arg1* and *Retnla*, whereas other cell types upregulated *Socs1* and *Socs3*, which are downstream of the TAM receptors (Rothlin et al. 2007).

**Figure 2:**
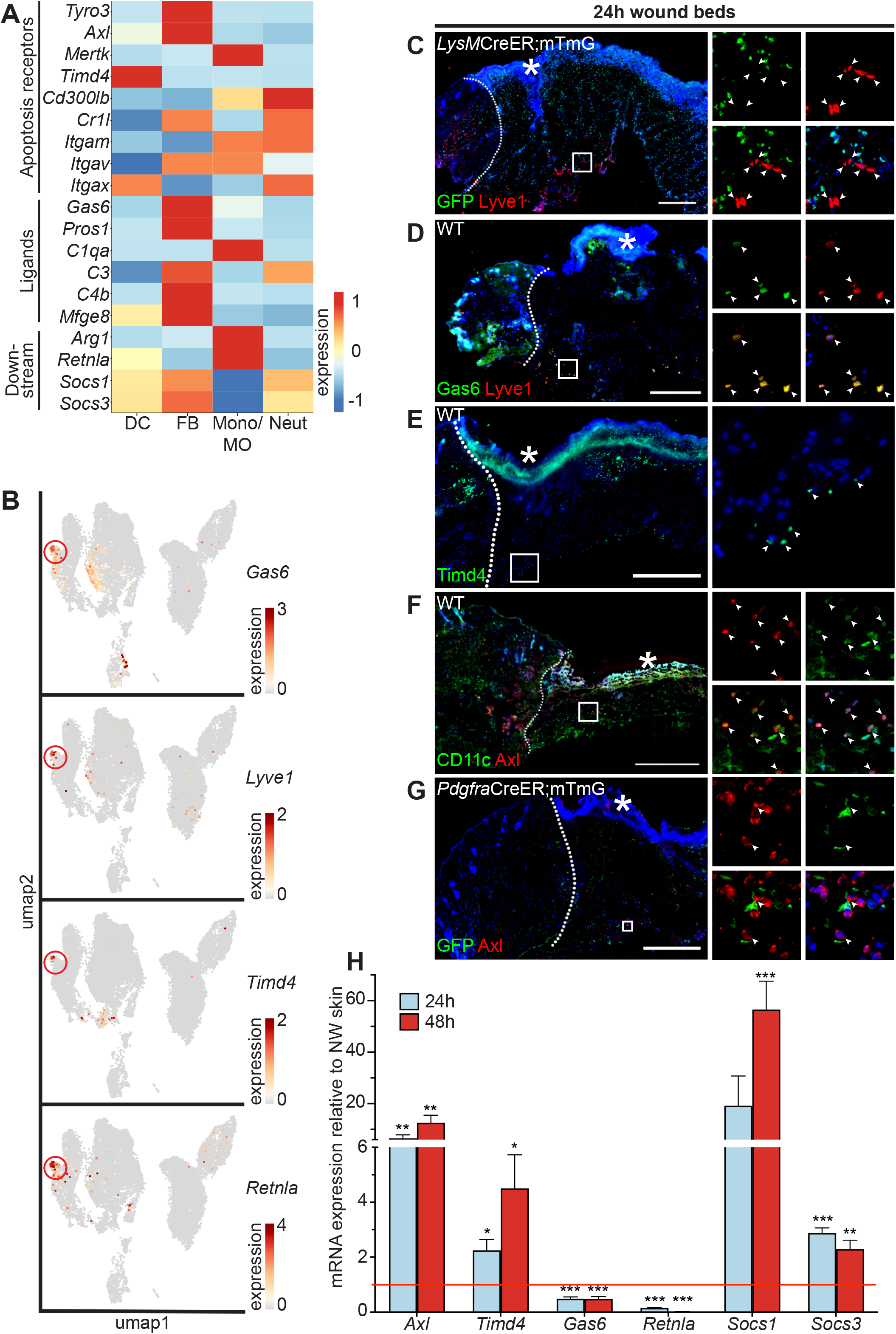
Apoptosis detection genes are highly expressed in the wound bed. A. Heatmap of differentially expressed efferocytosis pathway genes in 24h and 48h wound beds. B. Feature plots showing expression of *Gas6, Lyve1, Timd4*, and *Retnla* with *Lyve1*^+^ region highlighted. C. Immunostaining for GFP (green) and Lyve1 (red) in LysMCreERmTmG wound bed 24h after injury. Arrows indicate Lyve1^+^ cells. D. Immunostaining for Gas6 (green) and Lyve1 (red) in wild-type (WT) wound bed 24h after injury. Arrows indicate double-positive Gas6^+^ Lyve1^+^ cells. E. Immunostaining for Timd4 (green) in WT wound bed 24h after injury. Arrows indicate Timd4^+^ cells. F. Immunostaining for CD11c (green) and Axl (red) in wound bed 24h after injury. Arrows indicate double-positive CD11c^+^ Axl^+^ cells. G. Immunostaining for GFP (green) and Axl (red) in PdgfraCreERmTmG wound bed 24h after injury. Arrows indicate double-positive GFP^+^ Axl^+^ cells. H. mRNA expression of genes relative to nonwounded (NW) control at 24h and 48h after injury. Red line indicates normalized control mRNA levels. Error bars indicate mean +/− SEM, unpaired T-test, *p < 0.05, **p < 0.01, ***p < 0.001. * indicates scab. Scale bars = 500µm.

We noted that Axl’s ligand, *Gas6*, and several other genes involved in efferocytosis were expressed predominantly in the fibroblast cluster, but were also lowly expressed in the monocyte/macrophage cluster (**Fig. 2A**). Examining the UMAP plot to determine the heterogeneity of efferocytosis gene expression within individual cell types, we observed that *Gas6* was highly expressed by a specific subset of monocyte/macrophage cells. These cells also overexpressed the resident macrophage marker *Lyve1*, the apoptosis receptor *Timd4*, and *Retnla*, a downstream factor of efferocytosis (**Fig. 2B**), indicating that they may play a role in apoptosis detection and response in wound healing. *Lyve1* has been identified as a marker for resident macrophages, which are distinct from the majority of wound macrophages that differentiate from bone marrow derived-monocytes and are recruited to the wound after injury (Lim et al. 2018, Wang et al. 2020). To confirm the presence of Lyve1^+^ resident cells *in vivo*, we used a *LysM*CreER;mTmG mouse model, in which myeloid cells in the bone marrow can be induced to express GFP prior to injury, such that any GFP^+^ cells observed in the wound bed are interpreted as newly recruited to the site of injury, while GFP^-^ cells are interpreted to be resident to the skin. We observed that Lyve1 was expressed at the protein level both in wound beds (**Fig. 2C**) and adjacent to the wound (**Fig. S2A)** with immunofluorescence staining, and confirmed that these cells were resident rather than recruited to the wound bed after injury, since they did not express GFP. Further, we confirmed that Lyve1^+^ cells co-expressed Gas6 protein (**Fig. 2D**) and that Timd4 protein (**Fig. 2E**) was expressed in wound beds.

The TAM receptor *Axl* was uniquely expressed in the single-cell dataset by both dendritic cells and fibroblasts (**Fig. 2A**). Using immunofluorescence staining, we confirmed Axl protein expression in 24h wound beds, and that Axl protein colocalized with markers for both dendritic cells (CD11c) (**Fig. 2F**) and fibroblasts (Pdgfra-GFP) (**Fig. 2G**), in which the elongated processes of the fibroblasts are GFP^+^ while the Axl stain is centralized. We also analyzed mRNA expression of apoptosis-related genes by qPCR to determine their expression in wound beds compared to naive skin (**Figs. 2H and S2B**). Efferocytosis receptor mRNAs were significantly upregulated in the wound bed compared to naive skin, while the ligand *Gas6* was significantly downregulated. The downstream factor *Retnla* was also significantly downregulated in the wound bed, while *Socs1* and *Socs3* were significantly upregulated. Taken together, these data indicate that multiple efferocytosis pathways are involved in the early inflammatory stage of wound healing.

### Cell death signaling in human diabetic & nondiabetic wounds

We next set out to explore whether these apoptosis and efferocytosis-related pathways were also relevant in pathological states associated with dysregulated wound healing, such as diabetes. Previous studies have indicated that apoptosis is increased in diabetic wounds, including an elevation in apoptotic lymphocytes (Arya 2014). Additional studies have shown that hypoxic environments (such as those found in diabetic wounds) increase macrophage efferocytosis (Wang 2023). We recently performed scRNA-seq analysis of cells derived from foot wounds of healthy and diabetic human patients (to be published elsewhere). Thus, we determined whether transcriptional changes in apoptosis and efferocytosis signaling were present and in which cell types. Strikingly, diabetic foot wounds upregulated expression of efferocytosis receptors, ligands, and downstream genes associated with efferocytosis signalling compared to nondiabetic samples (**Fig. 3A**). In particular, mesenchymal cells (fibroblasts and pericytes), T cells, and lymphatic vessels expressed overall higher levels of mRNAs associated with efferocytosis signaling. Similarly, diabetic foot ulcers upregulated mRNAs involved in apoptotic signaling pathways compared to non-diabetic foot ulcers (**Fig. 3B**). Macrophages in diabetic wounds upregulated more apoptosis related mRNAs compared to macrophages in control wounds. These data further indicate an impact of diabetes on induction of efferocytosis and apoptotic genes in foot wound repair.

**Figure 3:**
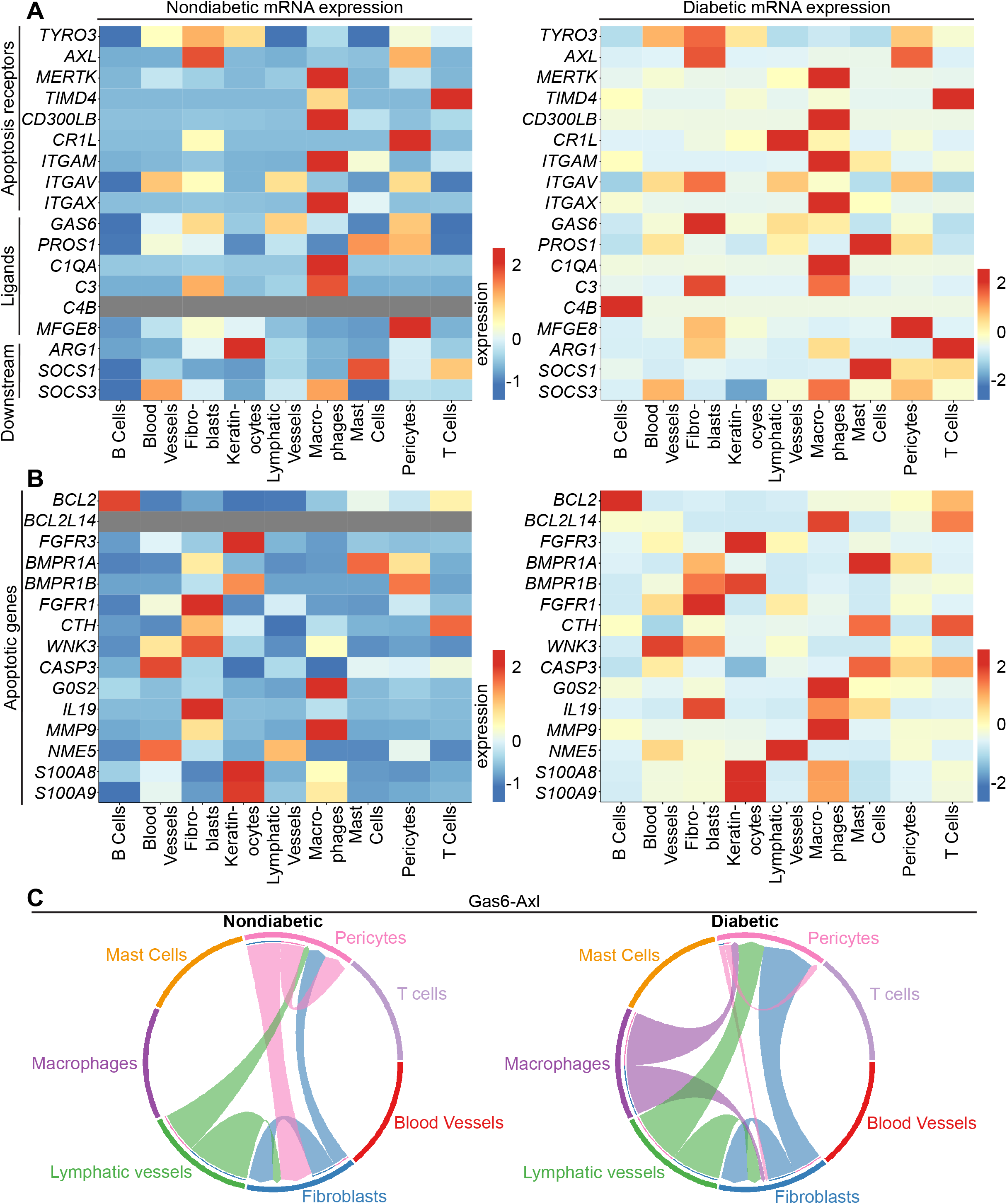
Human diabetic wounds have increased apoptosis signaling expression compared to nondiabetic wounds. A. Heatmap of differentially expressed genes related to apoptosis detection in nondiabetic and diabetic wound beds. B. Heatmap of differentially expressed genes related to apoptosis in nondiabetic and diabetic wound beds. C. CellChat Chord diagrams showing Gas6/Axl communication in nondiabetic and diabetic wound beds.

Interestingly, analysis of these data with CellChat, which quantitatively infers intercellular signaling networks from scRNA-seq data (Jin et al. 2021), revealed altered signaling between *Axl* and its ligand *Gas6* between the two groups. In non-diabetic patient wounds, *Gas6* expression by pericytes and lymphatic vessels stimulated signaling via *Axl* receptors on fibroblasts. However, in diabetic patient wounds, *Gas6* expression by macrophages activated *Axl* receptors on fibroblasts and pericytes and *Gas6* stimulation of Axl receptors by pericytes was reduced, while fibroblasts and lymphatic vessels displayed stronger pathway activation in fibroblasts and pericytes in diabetic foot wounds compared to control foot wounds (**Fig. 3C**). Taken together, these data indicated that diabetic wounds activate and modulate *Gas6-Axl* signaling.

### TLR3 stimulation is sufficient, but not necessary, for *Axl* upregulation in skin

Next, we were interested in examining the molecular mechanisms that induce *Axl* mRNA expression after injury. Prior work showed that *Axl* expression was induced by toll-like receptor 3 (TLR3) stimulation (Rothlin et al 2007) and that TLR3 is essential for skin wound repair (Lin et al. 2011). Thus, we experimentally tested the role of TLR3 signaling in *Axl* expression in the skin. scRNA-seq of early wounds revealed that *TLR* mRNAs are highly expressed in neutrophils and macrophages with a few expressed in dendritic cells and fibroblasts (**Fig. 4A**). Interestingly, *TLR3* is unique among the TLRs in that it is expressed by both dendritic cells and fibroblasts, which also express high levels of *Axl* in the single-cell dataset (**Fig. 2A**). To determine if TLR3 stimulation upregulates *Axl* expression in the skin, we injected the synthetic double-stranded RNA polyinosinic:polycytidylic acid (poly(I:C)) or a PBS control in naive mouse back skin of either wild-type (WT) or TLR3 knockout (KO) mice. We collected the injection site and surrounding area after 2 hours and processed the skin samples for qPCR and immunostaining (**Fig. 4B**). We first analyzed cytokine mRNA expression, a target of TLR3 signaling that promotes inflammation (Rothlin et al. 2007). While several inflammatory cytokines were not induced in skin injected with poly(I:C) (**Fig. S3A**), interferon (IFN)-β (*Ifnb)* was upregulated in WT mice injected with poly(I:C) but not in the skin of TLR3 KO mice (**Fig. 4C**), confirming the specificity of poly(I:C) for activation of TLR3 in the skin (Alexopoulou et al. 2001). Axl protein (**Fig. 4D**) and mRNA (**Fig. 4E**) were induced in naive skin upon poly(I:C) injection, and *Axl* mRNA induction and protein expression were abrogated in skin of TLR3 KO mice (**Figs. 4E and S3B**). Axl protein expression around the dermal injection site was quantified via corrected total fluorescence (**Fig. 4F**), confirming these results. Taken together, these data indicate that TLR3 signaling is sufficient to activate *Axl* expression in naive skin.

**Figure 4:**
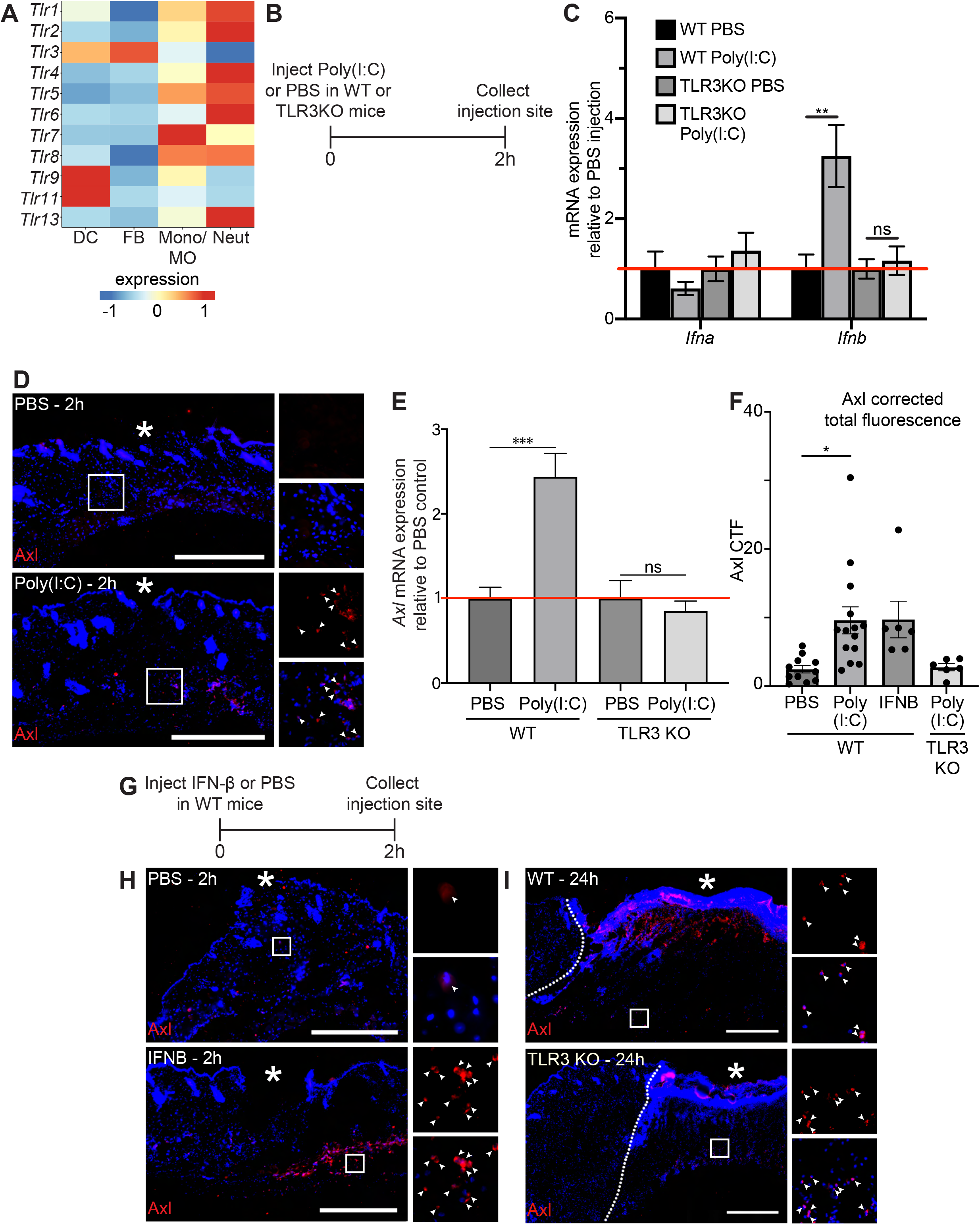
TLR3 signaling is sufficient to upregulate Axl in naive skin, but TLR3 is not required for Axl expression in the wound bed. A. Heatmap of differentially expressed TLR genes in 24h and 48h wound beds. B. Schematic of injection and collection protocol. C. mRNA expression of interferon genes relative to respective PBS injection. Red line indicates normalized control mRNA levels. Error bars indicate mean +/− SEM, unpaired T-test, **p < 0.01. ns, nonsignificant. D. Immunostaining for Axl (red) in naive back skin injected with PBS or Poly(I:C). Arrows indicate Axl^+^ cells. * indicates injection site. E. mRNA expression of *Axl* relative to respective PBS injection control 2h after injection. Red line indicates normalized control mRNA levels. Error bars indicate mean +/− SEM, unpaired T-test, ***p < 0.001. ns, nonsignificant. F. Quantification of corrected total fluorescence for Axl immunostaining in a 1mm square containing the injection site. Error bars indicate mean +/− SEM, one way ANOVA with multiple comparisons, *p < 0.05. G. Schematic of injection and collection protocol. H. Immunostaining for Axl (red) in naive back skin injected with PBS or IFNB. Arrows indicate Axl^+^ cells. * indicates injection site. I. Immunostaining for Axl (red) in wound beds 24h after injury in WT or TLR3KO mice. Arrows indicate Axl^+^ cells. * indicates scab. Scale bars = 500µm.

Since *Ifnb* was elevated by TLR3 signaling in naive skin, we determined whether injecting recombinant IFN-β intradermally into naive mouse back skin was sufficient to induce Axl protein expression (**Fig. 4G**). Two hours after injection, Axl protein was detected by immunostaining in skin injected with IFN-β but not control skin (**Figs. 4H,F**). Next, to determine if TLR3 activation was necessary for Axl expression in skin wounds, we analyzed Axl expression in wound beds of TLR3 KO mice. In contrast to our previous results in naive skin, Axl protein expression was stimulated in TLR3 KO wound beds similar to WT mice (**Fig. 4I**). Thus, these data suggest that while TLR3 is sufficient to drive Axl expression in naive skin, additional mechanisms drive Axl upregulation within skin wounds in its absence.

### Axl is required for skin wound healing

Based on our data showing strong upregulation of Axl mRNA and protein expression in wound beds, even in the absence of TLR3, we hypothesized that Axl may play a role in wound healing. To examine whether Axl signaling was required for wound repair, we intraperitoneally injected mice 3 hours prior to injury with either a control IgG antibody or anti-Axl function blocking antibody (Ab), which binds to Axl’s extracellular domain and blocks Axl-mediated viral infection (Retallack et al. 2016), and has been shown to inhibit Axl activity *in vitro* (**Fig. 5A**). Since Axl activity upregulates *Axl* mRNA expression in a positive feedback loop (Brand et al. 2014), we analyzed *Axl* mRNA expression. While we saw a non-significant downregulation of *Axl* mRNA at 1 day (D) post injury, *Axl* mRNA was significantly downregulated in skin wounds treated with anti-Axl Ab on day 5 compared to control IgG Ab injected wounds (**Fig. 5B**), indicating that the anti-Axl Ab treatment reduced Axl signaling in skin wounds. When we examined the effect of Axl inhibition on apoptotic cell clearance in skin wounds using TUNEL staining of skin sections derived from mouse wounds 1 day after injury, TUNEL^+^ cells were present but rare in both anti-Axl Ab and IgG Ab treated wounds (**Fig. S4A**), suggesting that additional efferocytosis mechanisms can clear apoptotic cells in early skin wounds despite Axl inhibition.

**Figure 5:**
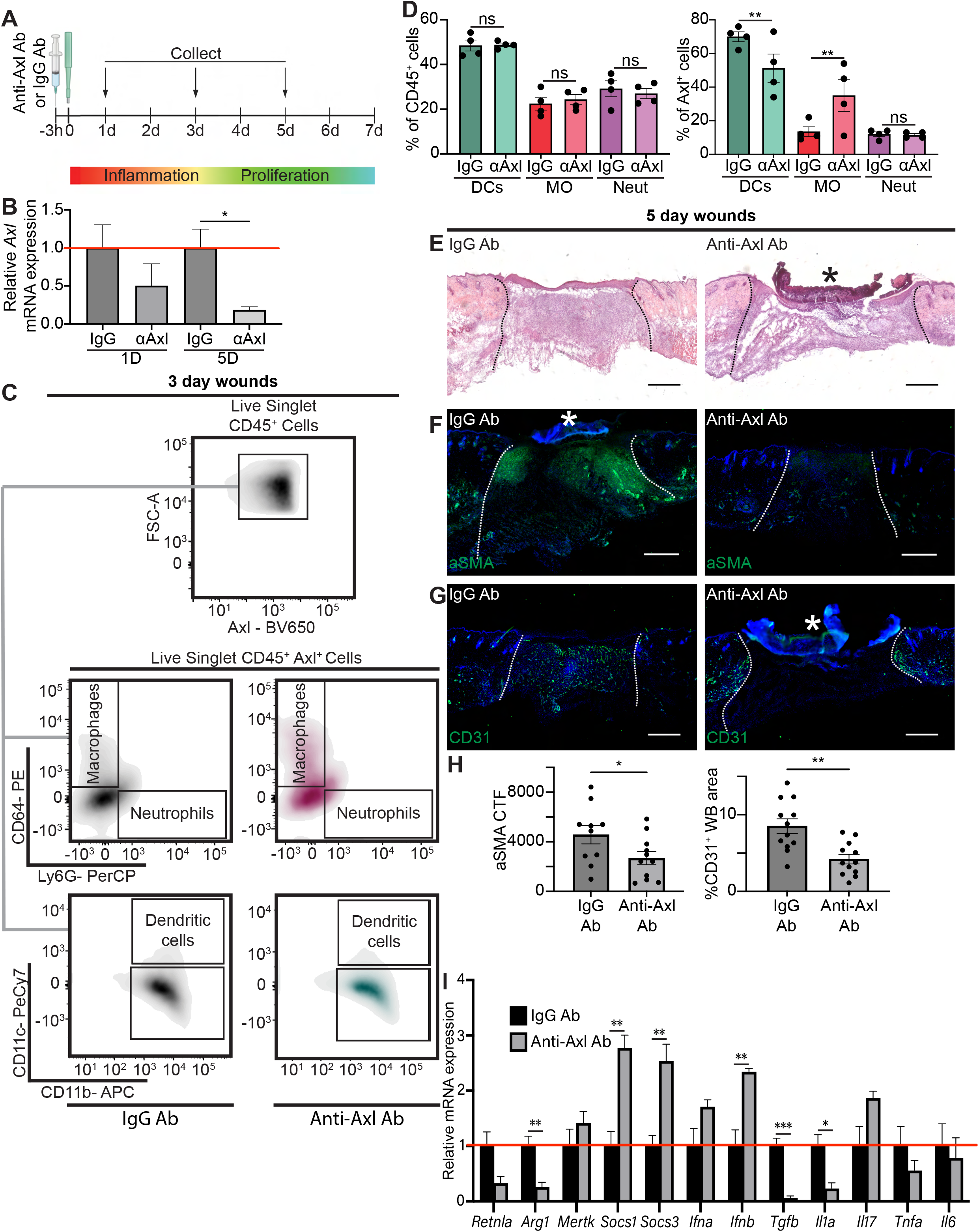
Axl antibody inhibition results in defects to wound repair and changes to inflammation. A. Schematic of experimental design. *B. Axl* mRNA expression normalized to respective IgG Ab control. Error bars indicate mean +/− SEM, one way ANOVA with multiple comparisons, *p < 0.05. C. Representative flow cytometry gates used to analyze cells isolated from wounds 3 days after injection and injury. Live singlet CD45^+^ cells were identified as macrophages, neutrophils, or dendritic cells via fluorescent antibody staining. D. **Left:** Quantification of CD45^+^ cells by cell type in anti-Axl Ab or IgG Ab treated wound beds 3 days after injury. Error bars indicate mean +/− SEM, one way ANOVA with multiple comparisons, ns, nonsignificant. **Right:** Quantification of Axl^+^ cells by cell type in anti-Axl Ab or IgG Ab treated wound beds 3 days after injury. Error bars indicate mean +/− SEM, two way ANOVA with multiple comparisons, **p < 0.01. ns, nonsignificant. E. H&E staining of wound beds 5 days after antibody injection and injury. * indicates scab. F. Immunostaining for aSMA (green) in wound beds 5 days after antibody injection and injury. * indicates scab G. Immunostaining for CD31 (green) in wound beds 5 days after antibody injection and injury. * indicates scab. H. **Left:** Quantification of aSMA corrected total fluorescence. Error bars indicate mean +/− SEM, unpaired T-test, *p < 0.05. **Right:** Quantification of CD31^+^ pixels in wound bed. Error bars indicate mean +/− SEM, unpaired T-test, **p < 0.01. I. mRNA expression of genes relative to IgG Ab control at 5 days after injury. Red line indicates normalized control mRNA levels. Error bars indicate mean +/− SEM, one way ANOVA with multiple comparisons, *p < 0.05, **p < 0.01, ***p < 0.001. Scale bars = 500µm

To characterize inflammation at 3 days after injury, we used flow cytometry to quantify the immune cells present in the wound bed (**Figs. 5C-D and S4B**). Interestingly, no significant change was observed in the proportion of dendritic cells, neutrophils, or macrophages that were present in wound beds of IgG or anti-Axl Ab treated wound beds. However, the proportion of cell types expressing Axl was altered, with more Axl^+^ macrophages and fewer Axl^+^ DCs present in the anti-Axl Ab treated group compared to the IgG Ab control. Despite the similar immune cell numbers in wounds in which Axl was inhibited, we did detect nonsignificantly increased expression of the inflammatory cytokines *Il1a* and *Il6* 1 day after injury (**Fig. S4C**), potentially indicating a trend towards greater inflammatory signaling in the anti-Axl Ab treated wound environment compared to the IgG Ab treated wounds.

Despite the relatively normal inflammatory cell numbers at day 3 after injury, wounds treated with anti-Axl Ab at 5 days post-injury exhibited observable healing defects compared to wounds treated with control IgG Ab, including a lack of granulation tissue and larger scab visible with H&E staining (**Fig. 5E**). We also observed that upon Axl inhibition, fibroblast repopulation was significantly reduced (**Figs. 5F and 5H**). Revascularization was also defective; Axl inhibition significantly reduced the CD31^+^ area of wound beds (**Figs. 5G-H**). While more anti-Axl Ab treated wound beds (8/12) failed to fully re-epithelialize than IgG Ab treated wounds (5/12), there was no significant difference in wound closure as indicated by ITGA6 staining between the two treatments (**Fig S4D**).

We found significant changes in gene expression at 5 days after injury, including a significant downregulation of *Arg1, Tgfb*, and *Il1a*, and significant upregulation of *Ifnb, Socs1*, and *Socs3* upon Axl inhibition (**Figs. 5I**). Taken together, these changes suggest that Axl inhibition impairs proper healing, potentially by altering the signaling response in immune cells, resulting in significant defects to revascularization and fibroblast repopulation.

### Timd4 function is required for normal skin repair

To further examine the role of efferocytosis receptors in skin wound repair, we abrogated Timd4 function prior to injury by intraperitoneally injecting a function blocking anti-Timd4 Ab, which effectively blocks efferocytosis in an atherosclerosis model (Foks et al. 2016). Strikingly, we found that TUNEL^+^ apoptotic cells were significantly more prevalent in 3 and 5 day wound beds treated with anti-Timd4 Ab compared to IgG Ab treated wound beds, (**Fig. 6A-B**), indicating a defect to efferocytosis.

**Figure 6:**
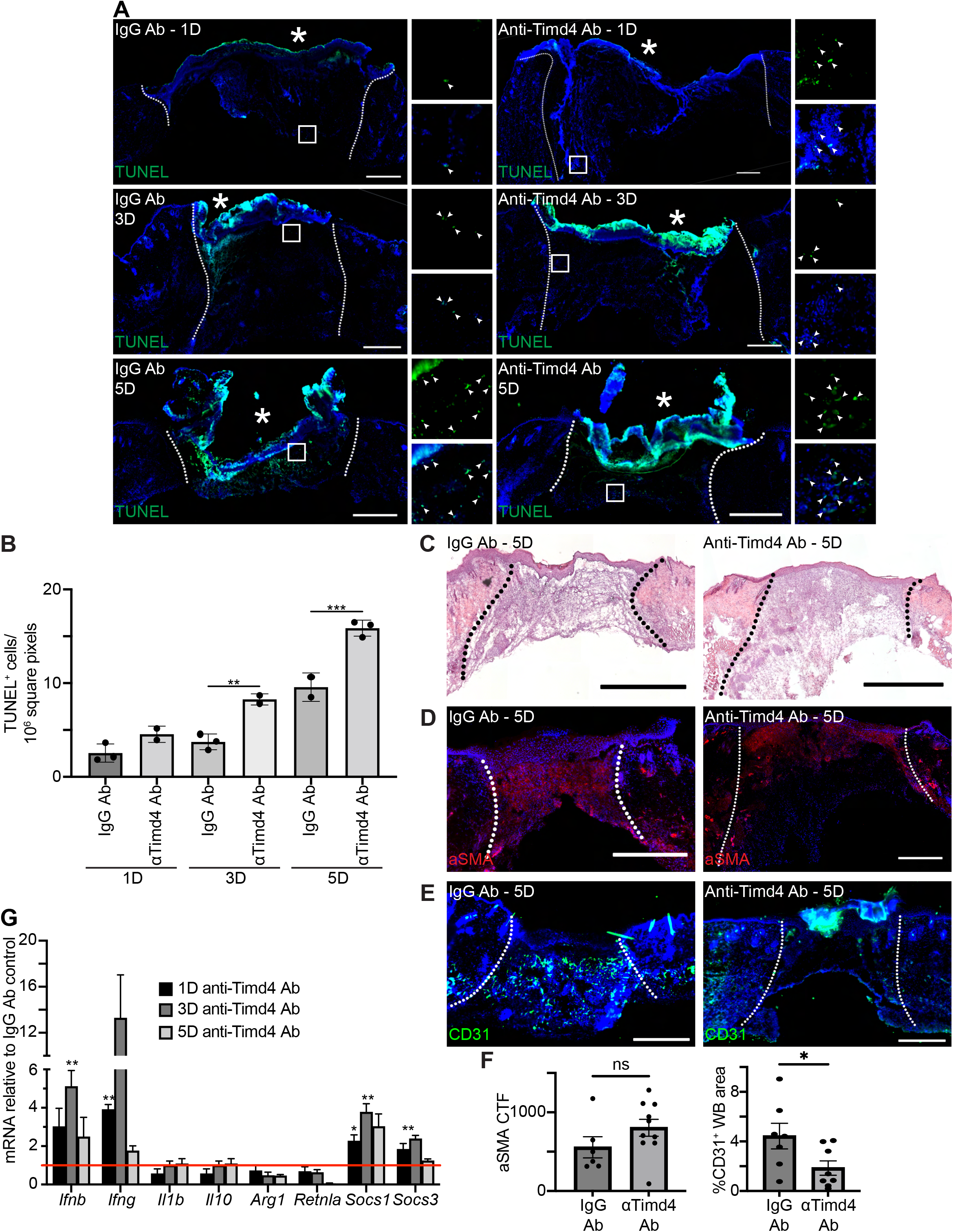
Timd4 function is required for efferocytosis during skin repair. A. Immunostaining for TUNEL (green) in wound beds injected with anti-Timd4 Ab or IgG Ab control 1, 3, or 5 days after injury. Arrows indicate TUNEL^+^ cells. * indicates scab. B. Quantification of TUNEL^+^ cells per 10^6^ square pixels. Error bars indicate mean +/− SEM, one way ANOVA, **p < 0.01, ***p < 0.001. C. H&E staining of wound beds 5 days after antibody injection and injury. D. Immunostaining for aSMA (red) in wound beds 5 days after antibody injection and injury. E. Immunostaining for CD31 (green) in wound beds 5 days after antibody injection and injury. F. **Left:** Quantification of aSMA corrected total fluorescence. Error bars indicate mean +/− SEM. ns, nonsignificant **Right:** Quantification of CD31^+^ pixels in wound bed. Error bars indicate mean +/− SEM, unpaired T-test, *p < 0.05. G. mRNA expression of genes relative to respective IgG Ab control. Red line indicates normalized control mRNA levels. Error bars indicate mean +/− SEM, unpaired T-test, *p < 0.01, **p < 0.01. Scale bars = 500µm.

Unlike inhibition of Axl, inhibition of Timd4 did not lead to obvious defects in overall healing as indicated by H&E staining of 5 day wounds (**Fig. 6C**), though we did observe qualitatively that the anti-Timd4 Ab treated wound beds appeared more fragile than the IgG Ab treated control wounds. Immunostaining of wound sections with antibodies against aSMA (**Figs. 6D and 6F**) and ITGA6 (**Figs. S5A-B**) did not indicate significant changes to either fibroblast repopulation or re-epithelialization, respectively. However, staining for CD31 (**Fig. 6E, 6F**) indicated a significant defect to revascularization in anti-Timd4 Ab treated wound beds compared to IgG Ab treated wounds.

Next, we examined mRNA expression profiles of inflammatory and efferocytosis signaling pathways in IgG Ab and anti-Timd4 Ab treated wound beds (**Fig. 6G**). The inflammatory cytokines *Ifng* and *Ifnb* were significantly upregulated at day 1 or 3, respectively, in Timd4 inhibited wounds compared to control wounds, suggesting altered inflammatory signaling. Interestingly, at day 1 and/or day 3, *Socs1* and *Socs3* were also significantly upregulated, similar to the gene expression pattern observed after Axl inhibition (**Fig. 5I**). Taken together, these results indicate that Timd4 activity is required for normal efferocytosis, inflammation gene expression, and revascularization after injury.

## Discussion

Here, we provide an atlas for the dynamic changes that occur in the early inflammatory stage of wound repair in the skin at the single-cell level. We found that apoptotic and efferocytosis pathways were upregulated in distinct cell types in mouse wounds and in diabetic foot wounds of human patients. Using functional inhibition studies in mouse wounds, our data show that efferocytosis receptors, Axl and Timd4, have differential effects on efferocytosis and wound repair.

We found that apoptotic genes were upregulated in most of the cell types present in early wounds, which likely allows the dynamic shifts in transcriptional profiles of inflammatory cells we observed in the first few days of tissue repair. Apoptotic cells are required for tail regeneration of *Xenopus laevis* (Tseng et al. 2007) and tissue regeneration of planaria (Hwang et al. 2004) and have been widely implicated in early wound repair (Greenhalgh 1998, Arya 2014). Mechanistically, apoptotic cells can release signaling factors including Wnts and PGE2 to promote proliferation of tissue resident cells and tissue repair in multiple species (Codispoti et al. 2019, Li et al. 2010, Fuchs et al. 2013). Detection of apoptotic cells has also been linked to inflammation and context-dependent integration of IL-4 signaling to activate an anti-inflammatory and tissue repair gene program (Bosurgi et al. 2017a).

Exposure of PtdSer on the outer leaflet of apoptotic cell membranes acts as a sensor for engulfment in evolutionarily conserved mechanisms (Rothlin & Ghosh 2020). Our data suggests that in skin wounds, apoptotic cells are rapidly detected and removed via efferocytosis, which is consistent with the large number of phagocytic neutrophils and macrophages in early wounds. We also found that mRNAs encoding efferocytosis receptors and signaling pathways are upregulated in the early skin wound beds and in diabetic foot wounds, and confirmed protein upregulation of the efferocytosis receptors Axl and Timd4 in murine wounds. Interestingly, efferocytosis receptors were upregulated in professional phagocytes as well as in fibroblasts, which also increased expression of a large number of efferocytosis ligands in both mouse wounds and diabetic foot ulcers (**Figs. 2 and 3**). These data resonate with recent data suggesting that fibroblasts can engulf apoptotic endothelial cells to alter their contractility, migration, and ECM production (Romana-Souza et al. 2021), and may indicate a direct role for fibroblasts in modulating the inflammatory milieu of early wounds through detection of apoptotic cells.

Growing evidence suggests that distinct efferocytosis receptors elicit cell- and tissue-specific responses. Axl and Mer exhibit differential ligand specificity and shedding up on activation (Zagórska et al. 2014). Furthermore, *Axl* null bone marrow-derived macrophages are unable to perform efferocytosis with TLR3 activation *in vitro*, which suggests that Axl plays a central role in efferocytosis in inflammatory environments (Bosurgi et al. 2017b). Here, we show that in early murine skin wounds, Axl inhibition did not impact efferocytosis in a detectable manner, but rather Axl expression shifted from dendritic cells to macrophages, suggesting possible compensatory mechanisms via other efferocytosis receptors. Furthermore, we did not detect differences in *Axl* expression in *TLR3* null and control wounds, further suggesting the inflammatory wound environment *in vivo* displays activation of *Axl* receptor expression independent of TLR3. Axl may function in skin wounds by interacting with other tyrosine kinase receptors, which has been shown for EGFR, MET, and PDGFR (Meyer et al. 2013). Interestingly, Axl/Gas6 signaling can also regulate tumorigenesis to support tumor cell survival (Linger et al. 2008), migration (Wilson et al. 2014), and angiogenesis (Li et al. 2009, Zhu et al. 2019). Future work exploring Axl’s function in a cell type dependent manner, including possibly directly activating angiogenesis in skin wounds, will decipher these possibilities.

Our data also implicate an important role for resident macrophages in regulating early wound repair mechanisms. In particular, we identified the expression of several efferocytosis receptors and ligands on Lyve1^+^ resident macrophages including Timd4. Timd4 is expressed on tissue resident macrophages in multiple tissues including in the peritoneal and resident cardiac macrophages. In the heart, Lyve1^+^ resident macrophages act in a cardioprotective manner during myocardial infarction (Dick et al. 2019) and are immunoregulatory and promote engraftment of cardiac allografts (Thornley et al. 2014). Interestingly, similar to our data, Timd4 promotes efferocytosis in the heart. During cardiac infarction, Timd4 also is required for T cell responses and support of regulatory T cells (Tregs) (Foks et al. 2016). In the skin, Timd4 is essential for allograft survival (Yeung et al. 2013), which has been proposed to act through DCs to support Tregs. Indeed, Timd4 is expressed in DCs in later stages of wound repair (Haensel et al. 2020). Our future studies will test the role of Timd4 on DCs and resident macrophages, T cell immunity, and how they impact wound repair phenotypes noted with Timd4 inhibition.

In summary, our data implicate apoptosis recognition receptors as an important regulator of skin wound healing. We find that early wound beds and human diabetic foot wounds significantly upregulate apoptotic genes and receptors that recognize apoptotic cells and that inhibition of multiple apoptotic receptors impairs wound repair. Given the importance of apoptosis in wound repair and the expression of distinct apoptotic receptors in different cells, skin wound healing is an excellent model to decipher the mechanisms by which distinct cells recognize and respond to interactions with apoptotic cells. Targeting these mechanisms may reveal therapeutic avenues that promote healing in chronic, non-healing wounds, particularly in diabetic patients.

## Methods

### Animals

Wild-type C57BL/6J mice (Strain #:000664), B6;129S1-Tlr3tm1Flv/J (TLR3 KO) mice (Strain #:005217), Lyz2tm1(cre/ERT2)Grtn/J (*LysM*CreER) mice (Strain #:031674), and B6.129(Cg)-Gt(ROSA)26Sortm4(ACTB-tdTomato,-EGFP)Luo/J (mTmG) mice (Strain #:007676) were purchased from The Jackson Laboratories. *Pdgfra*CreER mice were developed in the laboratory of B. Hogan (Duke University, Durham, NC). Mice were maintained through routine breeding in an Association for Assessment and Accreditation of Laboratory Animal Care (AALAC)-accredited animal facility at Yale University. Animals were maintained on a standard chow diet ad libitum (Harlan Laboratories, 2018S) in 12-hour light/dark cycling. Up to five injured mice were housed per cage. All experimental procedures were approved and in accordance with the Institutional Animal Care and Use Committee. For experiments using intraperitoneal (i.p.) tamoxifen administration, 100 μl of 30 mg/mL tamoxifen (Sigma Aldrich) in sesame oil was injected daily for three days prior to experiments.

### Human Subjects

Diabetic and non-diabetic adults with chronic foot ulcers that were undergoing skin wound debridement were consented to donate discarded tissue for this study (IRB approval # 1609018360). The diabetic foot ulcer specimens were obtained from 5 individuals diagnosed with Type 2 diabetes. The non-diabetic foot ulcer specimens were obtained from the same individual, 2 of which were from the same wound, one month apart from each other. This individual suffered from peripheral neuropathy, peripheral vascular disease, and chronic venous stasis, which all together might have contributed to the presence of chronic foot wounds.

### Wounding

7-9 week-old mice were wounded during the telogen phase of hair cycling. Mice were anesthetized using isoflurane and six full-thickness wounds, at least 4 mm apart, were made on shaved back skin using a 4 mm biopsy punch (Millitex). Animals were sacrificed at noted intervals after injury and wound beds were processed for subsequent analysis.

### Single-cell digestion of mouse wounds

Wound beds were digested for single-cell RNA sequencing analysis in a buffer of Roswell Park Memorial Institute (RPMI) medium with Glutamine (Gibco), Liberase Thermolysin Medium (TM) (Roche), DNase, N-2-hydroxyethylpiperazine-N-2-ethane sulfonic acid (HEPES) (Gibco), Sodium Pyruvate (Gibco), Non-essential amino acids (Gibco), and Antibiotic-Antimycotic (100X) (Gibco). Blood cells were removed with Ammonium-Chloride-Potassium (ACK) lysing buffer (Gibco). Cells were resuspended in Dulbecco’s Modified Eagle’s Medium (DMEM) (ATCC) with 0.1% Bovine Serum Albumin (BSA) for analysis.

### Human skin collection and processing

Skin wound specimens were collected at the clinical setting in PBS with Antibiotic-Antimycotic (100X) (Gibco), and transported to the lab on ice for processing. All specimens were processed within 3 hr. of collection. The specimen was cleaned by immersion with 10% Betadine, 70% ethanol, and PBS. Excess blood and subcutaneous fat were removed, and skin was mechanically minced before being digested in an enzyme cocktail consisting of dispase (Stemcell Technologies), collagenase I (Worthington), and collagenase II (Worthington) in 0.25% Trypsin-EDTA (Gibco). Blood cells were removed with Ammonium-Chloride-Potassium (ACK) lysing buffer (Gibco). Cells were resuspended in Dulbecco’s Modified Eagle’s Medium (DMEM) (ATCC) with 0.1% Bovine Serum Albumin (BSA) for analysis.

### Single-cell data of mouse samples

scRNA-seq data from 24 and 48 hour mouse wound beds were processed using the standard cellranger pipeline (10x Genomics). Downstream analysis was performed using the Scanpy package in Python (Wolf et al. 2018). Cells were filtered for quality control to avoid doublets and dead cells. Dimensionality reduction and downstream data visualization were completed using the Scanpy implementation of UMAP (McInnes et al. 2020) and the ShinyCell package in R (Ouyang et al. 2021), respectively. Data is presented as scaled log-normalized mRNA counts (i.e., expression).

We adapted the cell type annotation pipeline from Kumar et al. (Kumar et al. 2018) to label our sequencing data by broad cell type, as was done in Wasko et al. (Wasko et al. 2022). Differentially expressed genes (DEGs) across timepoints were calculated using the rank_genes_groups function from the Scanpy module in Python with default parameters. We then performed enrichment analysis on the top (logfoldchanges > 1.5) DEGs for each group using g:Profiler (Raudvere et al. 2019), with the Gene Ontology (GO) knowledgebase as the reference database.

Link to GitHub repository: https://github.com/khbridges/justynski-woundbed

### Single-cell data of human samples

Human scRNA-seq data from human samples were generated from 10X Genomics 3’-end single cell gene expression V2. Analysis was performed in Scanpy (Wolf et al. 2018). Cells were filtered for doublets and dead cells for downstream analysis. Batch correction was performed to integrate cells across samples using Scanorama (Hie et al. 2019). Gene expression was scaled, log-transformed, and normalized. Dimensionality reduction was done using PCA and UMAP in Scanpy (McInnes et al. 2020). Differentially expressed genes were calculated using Wilcoxon rank_genes_groups function from Scanpy. Cell-cell communication analysis and data visualization of circos plot were performed using CellChat (Jin et al. 2021). CellChat is superior to similar methods because its algorithm accounts for the roles of both signaling cofactors and protein-protein signaling in its predictions of ligand-receptor interaction (Bridges & Miller-Jensen 2022). CellChat is available as an open-source software package in R.

### Staining and imaging

Mouse skin and wound beds were embedded in optimum cutting temperature (OCT) compound (VWR) and wound beds were sectioned through their entirety to identify the center. 7 µm or 14 µm cryosections were processed as previously described (Shook et al. 2016) and stained with antibodies listed below or hematoxylin and eosin. Composite images were acquired using the tiles module on a Zeiss AxioImager M1 (Zeiss) equipped with an Orca camera (Hamamatsu).

### Quantitative real-time PCR

Whole wound bed samples were digested using TRIzol LS (Invitrogen). RNA was extracted from the aqueous phase using the RNeasy Plus Mini Kit (Qiagen). cDNA was generated using equal amounts of total RNA with the Superscript III First Strand Synthesis Kit (Invitrogen) per manufacturer instructions. All quantitative real-time PCR was performed using SYBR green on a LightCycler 480 (Roche). Primers for specific genes are listed below. Results were normalized to β-actin as previously described.

### Injections

Mice were injected intradermally with 10 uL PBS, 50 ug/mL Poly(I:C) (Invivogen), or 5ug/mL IFNB (R&D) with 0.5% BSA. The injection site was isolated using a 6mm biopsy punch 2 hours after injection and processed for staining.

To inhibit signaling pathways *in vivo*, mice were injected intraperitoneally with 25ug/100uL anti-Axl antibody (R&D), 200ug/100uL anti-Timd4 (BioXCell), or equivalent unit IgG control (R&D) in PBS three hours before wounding.

### Flow cytometry

Mouse wound beds were dissected and digested into single cells using Liberase TM (Roche) and cells were suspended in fluorescence-activated single cell sorting (FACS) staining buffer (0.05% BSA in DMEM). Digested tissue was filtered with a 70 μm and 40 μm cell strainer prior to centrifugation. Cell suspensions were stained with antibodies for 30 minutes on ice. Dendritic cells were defined as CD11b^+^ Cd11c^+^ cells; Macrophages were defined as CD11b^+^ CD11c^-^ CD64^+^ Ly6G^-^ cells; Neutrophils were defined as CD11b^+^ CD11c^-^ CD64^-^ Ly6G^+^ cells. To exclude dead cells, Sytox Blue (Invitrogen, 1:1000) was added immediately before analysis or sorting using a FACS Aria III with FACS DiVA software (BD Biosciences).

To quantify cell death, mouse wound beds and naive back skin were digested as above and stained with an Annexin V Conjugates for Apoptosis Detection kit (Invitrogen). Propidium iodide (Invitrogen, 1:500) was added immediately before analysis using a BD FACS LSR Fortessa X20.

Flow cytometry analysis was performed using FlowJo Software (FlowJo).

### Image quantification

Histological quantification for each wound bed was conducted on multiple central sections for each wound bed when available. The percentage of the wound bed covered by ITGA6 staining (re-epithelialization) and corrected total fluorescence for Axl (in a 1mm square around the injection site) or aSMA (in the wound bed) were calculated using ImageJ software (National Institutes of Health, Bethesda, MD) as described previously (Schmidt and Horsley, 2013; Shook et al. 2018). Revascularization (CD31^+^) was calculated using Adobe Photoshop to measure the total pixels positive for antibody staining divided by the total number of pixels in wound beds. Cell death was quantified by counting clearly distinguished TUNEL^+^ nuclei and dividing by the total area of wound beds.

### Statistics

To determine significance between two groups, comparisons were made using Student’s t-test. Analyses across multiple groups were made using a one- or two-way ANOVA with Bonferroni’s post hoc using GraphPad Prism for Mac (GraphPad Software, La Jolla, CA) with significance set at p < 0.05.

### Antibodies

TUNEL kit: Click-iT™Plus TUNEL Assay for In Situ Apoptosis Detection, Alexa Fluor™488 dye Thermo Fisher C10617

Active Caspase-3: Human/Mouse Active Caspase-3 Antibody R&D AF835

Axl: AXL Polyclonal Antibody Bioss BS-5180R

CD31: Purified Rat Anti-Mouse CD31 BD Biosciences 550274

CD31: Anti-PECAM-1 Antibody, clone 2H8, Azide Free Millipore Sigma MAB1398Z

CD11c: Anti-CD11c antibody [N418] abcam ab33483

Gas6: GAS 6 Polyclonal Antibody, ALEXA FLUOR® 488 Conjugated Bioss BS-7549R-A488

GFP: Anti-GFP antibody abcam ab13970

ITGA6: Human/Mouse/Bovine Integrin alpha 6/CD49f Antibody R&D MAB13501

Lyve1: Anti-LYVE1 antibody - BSA and Azide free abcam ab14917

Timd4: TIM-4 Polyclonal Antibody Invitrogen PA5-116045

aSMA: Anti-alpha smooth muscle Actin antibody Abcam ab5694

### FACS antibodies

Axl - BV650 - 748032 - 1:200

Ly6G - PerCP - 127654 - 1:400

CD206 - AF488 - 141710 - 1:250

CD11c - Pe-Cy7 - 117318 - 1:500

CD40 - Pe-Cy5 - 124618 - 1:200

CD64 - PE - 139304 - 1:300

CD45 - APC-eFluor780 / APC-Cy7 - 47-0451-82 - 1:200

CD11b - APC - 101212 - 1:200

Sytox Blue - Invitrogen S34857

Annexin V Conjugates for Apoptosis Detection - Invitrogen A13201

Propidium Iodide - Invitrogen P3566

### Primers

Arg1 F: CATTGGCTTGCGAGACGTAGAC

Arg1 R: GCTGAAGGTCTCTTCCATCACC

Axl F: CGAGAGGTGACCTTGGAAC

Axl R: AGATGGTGGAGTGGCTGTC

Β-actin F:ATCAAGATCATTGCTCCTCCTGAG

B-actin R: CTGCTTGCTGATCCACATCTG

C3 F: CCAGCTCCCCATTAGCTCTG

C3 R: GCACTTGCCTCTTTAGGAAGTC

C4b F: ACTTCAGCAGCTTAGTCAGGG

C4b R: GTCCTTTGTTTCAGGGGACAG

Cd47 F: TGCGGTTCAGCTCAACTACTG

Cd47 R: GCTTTGCGCCTCCACATTAC

Cd300lb F: TGCAGGGTCCTCATCCGAT

Cd300lb R: TGTCCGTGTCATTTTGCCTGA

C1qa F: ATGGAGACCTCTCAGGGATG

C1qa R: ATACCAGTCCGGATGCCAGC

Cr1l F: ATGGAGGTCTCTTCTCGGAGT

Cr1l R: GGCCGAAGGCTACAAGGAG

Gas6 F:ATGAAGATCGCGGTAGCTGG

Gas6 R: CCAACTCCTCATGCACCCAT

Gla F: TCTGTGAGCTTGCGCTTTGT

Gla R: GCAGTCAAGGTTGCACATGAAA

Ifna F: CTTCCACAGGATACTGTGTACCT

Ifna R: TTCTGCTCTGACCACCCTCCC

Ifnb F: ATGAGTGGTGGTTGCAGGC

Ifnb R: TGACCTTTCAAATGCAGTAGATTCA

Ifng F: CAGGCCATCAGCAACAACATAAGC

Ifng R: ACCCCGAATCAGCAGCGACTC

Il10 F: GCCCAGAAATCAAGGAGCATT

Il10 R: TGCTCCACTGCCTTGCTCTTA

Il17 F: GTCGAGAAGATGCTGGTGGGTGTG

Il17 R: ACGTGGGGGTTTCTTAGGGGTCAG

Il6 F: AGCCCACCAAGAACGATAGTC

Il6 R: TTGTGAAGTAGGGAAGGCCG

Il1a F: TTGGTTAAATGACCTGCAACA

Il1a R: GAGCGCTCACGAACAGTTG

Il1b F: CTCATTGTGGCTGTGGAGAAG

Il1b R: ACACTAGCAGGTCGTCATCAT

Itgam F: TTCCTGGTGCCAGAAGCTGAA

Itgam R: CCCGTTGGTCGAACTCAGGA

Itgav F: CCGTGGACTTCTTCGAGCC

Itgav R: CTGTTGAATCAAACTCAATGGGC

Itgax F: CCCACCACTTCCTCCTGTAAC

Itgax R: AGCAATTGGGTCACAGGTTC

Mertk F: GGCTTTTGGCGTGACCATG

Mertk R: AGTTCATCCAAGCAGTCCTC

Mfge8 F: AGATGCGGGTATCAGGTGTGA

Mfge8 R: GGGGCTCAGAACATCCGTG

Pros1 F: TGGCAAGGAGACAGGTGTCAGT

Pros1 R: GAGCAGTGGTAACTTCCAGGAG

Retnla F: CCAATCCAGCTAACTATCCCTCC

Retnla R: CCAGTCAACGAGTAAGCACAG

Sirpa F: CCACGGGGAAGGAACTGAAG

Sirpa R: ACGTATTCTCCTGCGAAACTGTA

Socs1 F: CCGTGGGTCGCGAGAAC

Socs1 R: AACTCAGGTAGTCACGGAGTACCG

Socs3 F: TCCCATGCCGCTCACAG

Socs3 R: ACAGGACCAGTTCCAGGTAATTG

Tgfb F: ACTGTGGAAATCAACGGGATCA

Tgfb R: CTTCCAACCCAGGTCCTTCC

Timd4 F: AGCTTCTCCGTACAGATGGAA

Timd4 R: CCCACTGTCACCTCGATTGG

Tnfa F: TGTCTACTCCTCAGAGCCCC

Tnfa R: TGAGTCCTTGATGGTGGTGC

Tyro3 F: GAGGATGTCCTCATTCCAGAGC

Tyro3 R: CACTGCCACTTTCACGAAGGAG

## Acknowledgements

We would like to thank members of the Horsley and Miller-Jensen laboratory for their feedback and critical analysis of these data and manuscript. We would also like to thank the Yale Animal Resources Center (YARC) staff for animal husbandry and the Yale Science Building (YSB) Imaging core facility for confocal use. V.H. is funded by N.I.H-NIAMS R01s AR076938, AR0695505, AR075412. K.M-J. is funded by NIH U01-CA238728, R01-CA238728, and R01-GM123011.

## Author Contributions

Olivia Justynski: Conceptualization, Investigation, Visualization, Validation, and Writing – Original Draft, Review, and Editing; Kate Bridges: Formal analysis, Conceptualization, Visualization, and Writing – Original Draft; Will Krause: Investigation and Validation; Quan Phan: Formal analysis and Investigation; Teresa Sandoval-Schaefer: Investigation; Ryan Driskell: Supervision; Kathryn Miller-Jensen: Conceptualization, Visualization, Funding acquisition and Writing – Original Draft, Review, and Editing; Valerie Horsley: Conceptualization, Visualization, Funding acquisition, Supervision, and Writing – Original Draft, Review, and Editing.

## Inclusion and Diversity

One or more of the authors of this paper self-identifies as: 1) an underrepresented ethnic minority in their field of research or within their geographical location, 2) a gender minority in their field of research, and 3) a member of the LGBTQ+ community. While citing references scientifically relevant for this work, we also actively worked to promote gender balance in our reference list.

## Figure legends

**Figure S1: Dynamic cellular heterogeneity and evidence of apoptosis is observed in 24h and 48h wound beds by single-cell RNA sequencing**

A. UMAP plot of scRNA-seq data for cells from 24h and 48h wound beds in murine back skin annotated by cell identity.

B. Feature plots showing expression of two marker genes for each major cell type.

C. Number of cells identified by cell type at 24h and 48h.

D. Proportion of total cells represented by 24h or 48h for each major cell type.

E. Venn diagrams representing significantly upregulated genes shared by both timepoints or specific to 24h or 48h for each major cell type.

F. Immunostaining for TUNEL (green) in 48h wound bed. Arrows indicate TUNEL^+^ cells. * indicates scab. Scale bar = 500µm.

G. Quantification of live, apoptotic, and necrotic cells by flow cytometry using Annexin V and Propidium iodide (PI).

H. Representative FACS dot plots of gating strategy to identify live, apoptotic, and necrotic cells via Annexin V and PI staining. Apoptotic quadrant (AV^+^ PI^-^) is highlighted.

**Figure S2: Lyve1 and apoptosis receptors, ligands, and downstream factors are expressed in 24h and 48h wound beds**

A. Immunostaining for GFP (green) and Lyve1 (red) in LysMCreERmTmG wound bed-adjacent skin 24h after injury. Arrows indicate Lyve1^+^ cells, * indicates scab. Scale bar = 500µm.

B. mRNA expression of genes relative to NW control at 24h and 48h after injury. Red line indicates normalized control mRNA levels. Error bars indicate mean +/− SEM, unpaired T-test, *p < 0.05, **p < 0.01, ***p < 0.001.

**Figure S3: Poly(I:C) injection does not elicit Axl expression in TLR3 KO model**

A. mRNA expression of cytokine genes relative to respective PBS injection. Red line indicates normalized control mRNA levels. Error bars indicate mean +/− SEM.

B. Immunostaining for Axl (red) in naive back skin of TLR3KO mice injected with PBS or Poly(I:C). Arrows indicate Axl^+^ cells. * indicates injection site. Scale bar = 500µm.

**Figure S4: Analysis of anti-Axl antibody and IgG control-treated wound beds**

A. **Left:** Immunostaining for TUNEL (green) in wound beds injected with anti-Axl Ab or IgG Ab control 1 day after injury. Arrows indicate TUNEL^+^ cells. * indicates scab. Scale bars = 500µm.

**Right:** Quantification of TUNEL^+^ cells per 10^6^ square pixels. Error bars indicate mean +/− SEM.

B. Representative FACS dot plots of gating strategy to identify single cells, CD45^+^ immune cells, Axl^+^ cells, and immune cell populations.

C. mRNA expression of genes involved in TAM receptor signaling and cytokines relative to IgG Ab control 1 day after injury. Red line indicates normalized control mRNA levels. Error bars indicate mean +/− SEM.

D. **Left:** Immunostaining for ITGA6 (green) in wound beds 5 days after antibody injection and injury.

**Right:** Quantification of % re-epithelialization for 5 day wound beds. Error bars indicate mean +/− SEM, ns, nonsignificant.

**Figure S5: Re-epithelialization is not significantly altered in anti-Axl antibody and IgG control-treated wound beds**

A. Immunostaining for ITGA6 (green) in wound beds 5 days after antibody injection and injury.

B. Quantification of % re-epithelialization for 5 day wound beds. Error bars indicate mean +/− SEM.

