## Supplementary figures and images for "Apoptosis recognition receptors regulate skin tissue repair in mice"

### Figure S1

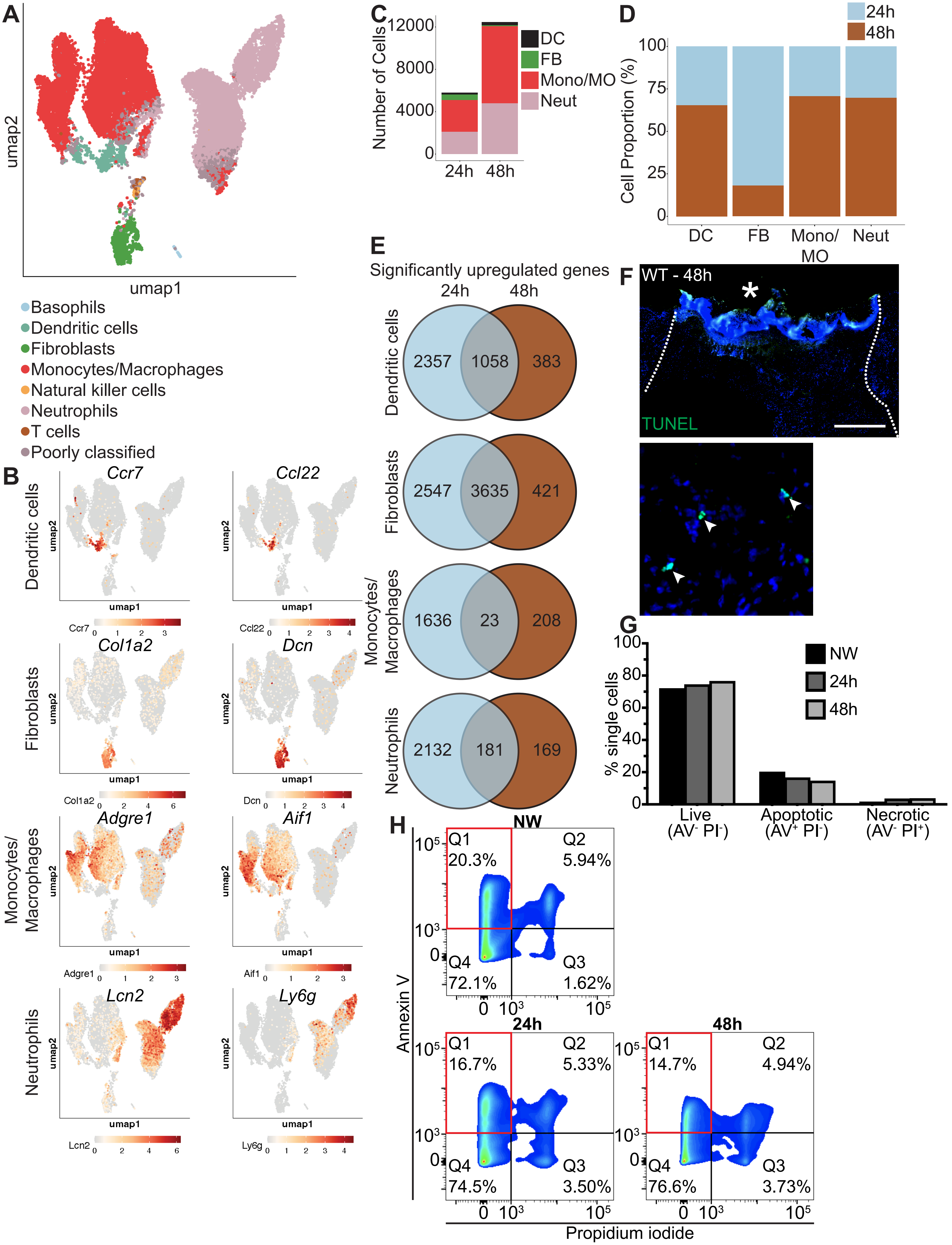

### Figure S2

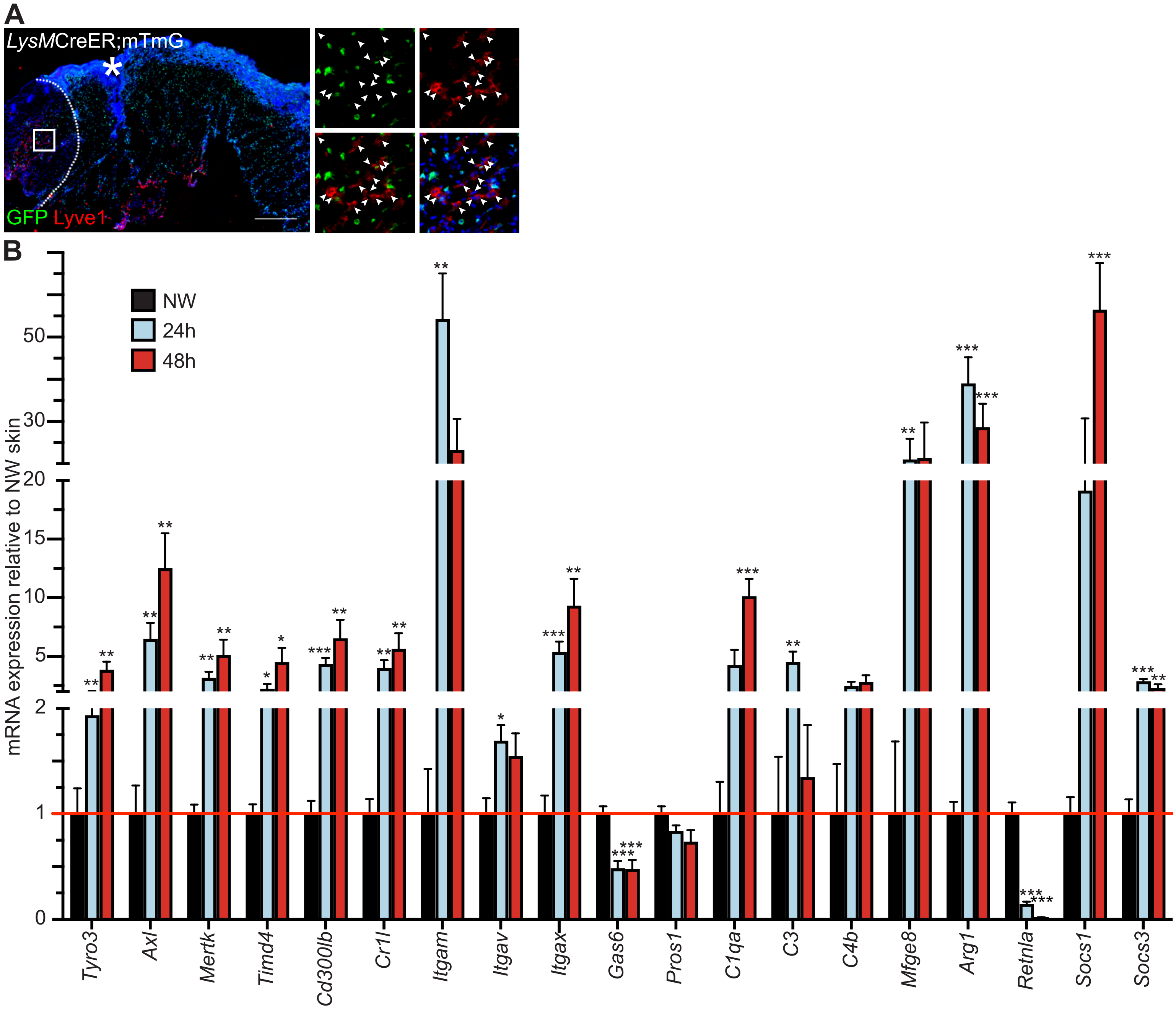

### Figure S3

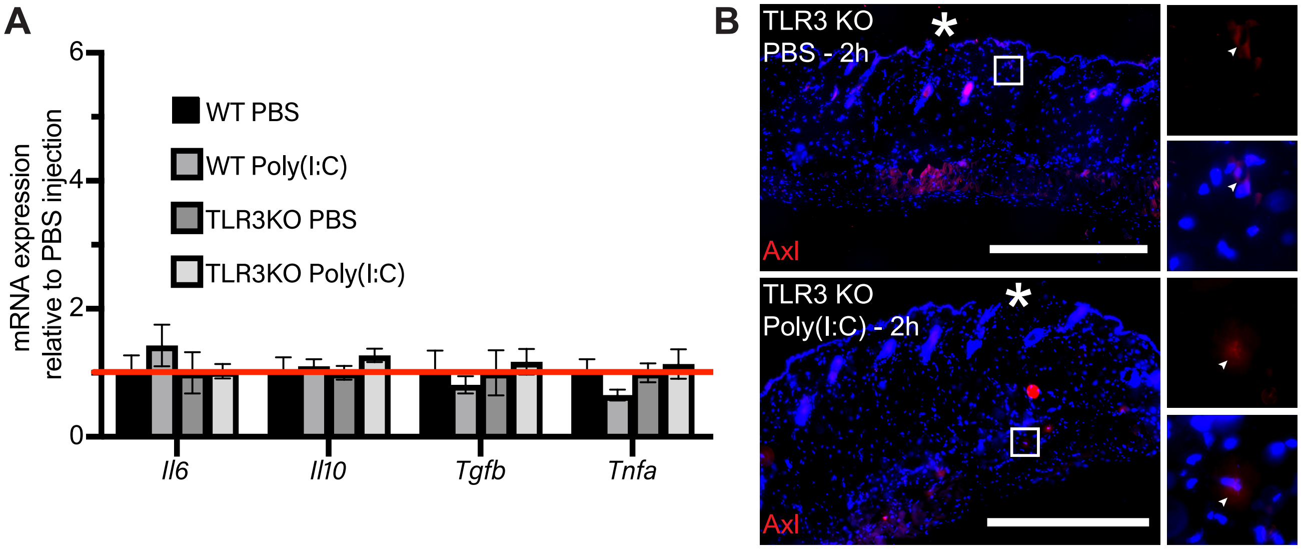

### Figure S4

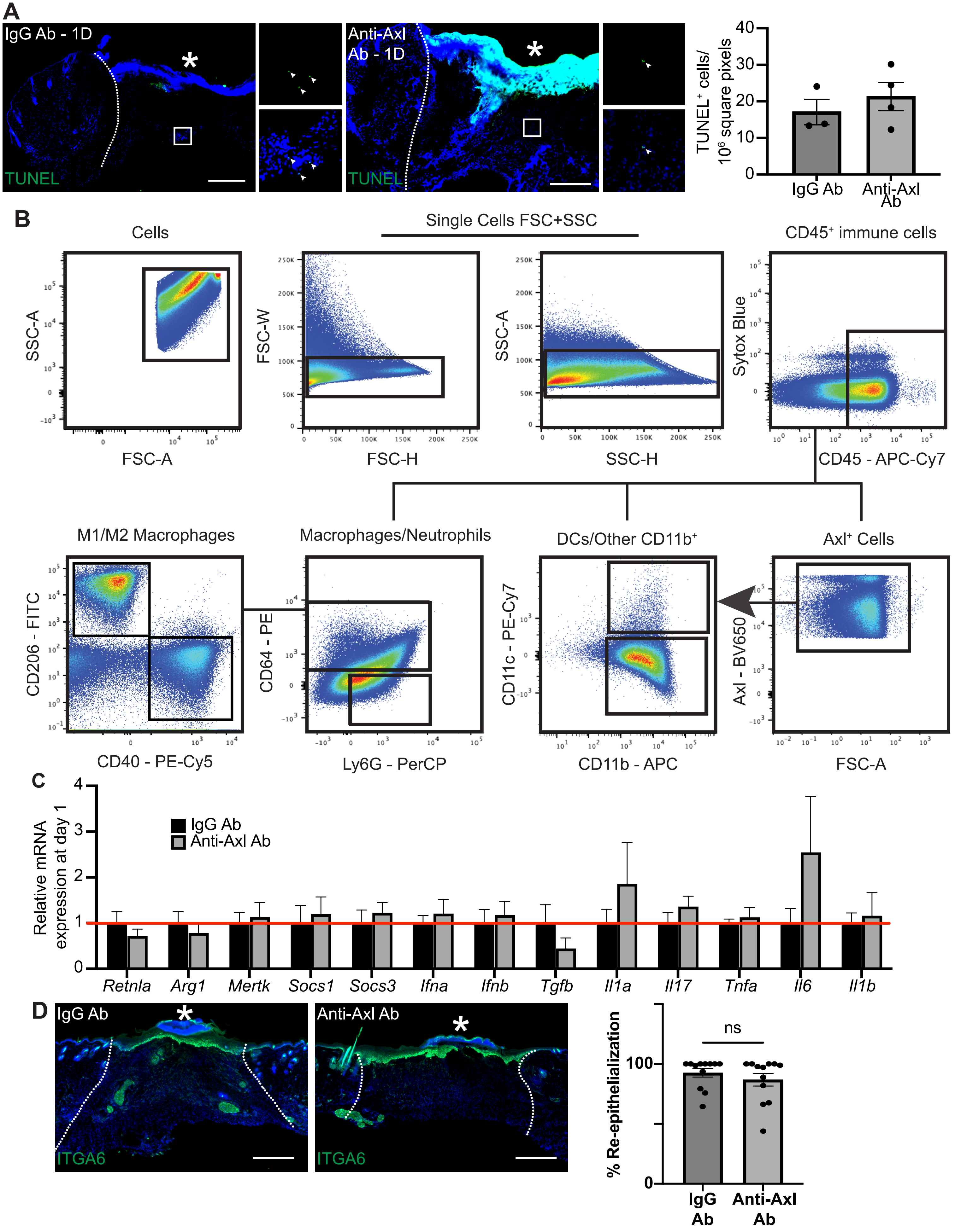

### Figure S5

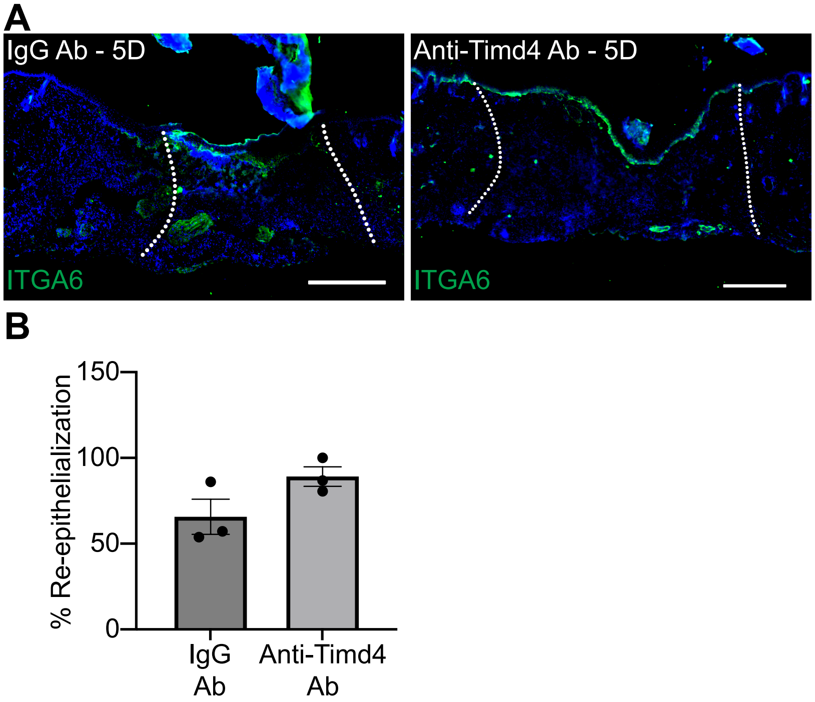
